# Accurate antibody loop structure prediction enables zero-shot design of target-specific antibodies

**DOI:** 10.1101/2024.08.23.609114

**Authors:** Yewon Bang, Yoon-Aa Choi, Jeonghyeon Gu, Sohee Kwon, Dohoon Lee, Eun Seo Lee, Myeong Sup Lee, Sangchoon Lee, Soyul Lee, Su Jung Lee, Jungsub Lim, Jin Young Maeng, Juno Nam, Jinsung Noh, Hyunjeong Oh, Sun-Young Park, Taeyong Park, Sumin Seo, Chaok Seok, Moo Young Song, Jonghun Won, Hyeonuk Woo, Jinsol Yang, Min Ji Yoon, Woong Bae, Jaehoon Kim, Dongjin Lee, Jaemyung Lee, Youhan Lee, Hasun Yu

## Abstract

Protein loops, characterized by their versatile structures with varying sizes and shapes, can recognize a wide range of targets with high specificity and affinity. The variable loops of the antibody complementarity-determining region (CDR) are particularly crucial for immune responses and therapeutic applications due to their effective target recognition capabilities. Accurate structure prediction of these antibody loops is essential for the efficient *in silico* design of target-binding antibodies for therapeutic or industrial use. However, predicting antibody loop structures is challenging due to the lack of evolutionary information from related proteins. Thus, a successful *ab initio* structure prediction method, which operates without structural templates or related sequences, is crucial for the effective design of antibody loop-mediated interactions. This study demonstrates that highly accurate antibody loop structure prediction enables the effective zero-shot design of target-binding antibody loops. The performance of loop design has been shown to depend on the accuracy of *ab initio* loop structure prediction, as tested with two versions of our design model. The high affinity, diversity, novelty, and specificity of the antibody loops designed with these new methods were validated experimentally on four target proteins.

## 1 Introduction

Structure-based *de novo* protein design has evolved from classical physics-based methods, such as RosettaDesign [1] to deep learning-based methods, such as RFdiffusion [2]. The significant improvement in protein design performance of RFdiffusion over RosettaDesign is primarily attributed to the enhanced accuracy in structure prediction, as demonstrated by advancements in RoseTTAFold and AlphaFold 2 [3, 4], as well as the integration of new protein generative models [5]. However, antibody binder design has remained challenging due to the low accuracy in predicting antibody loop-mediated complex structures. Recently, RFdiffusion-Ab has been reported for the *de novo* design of single-chain antibodies to several targets [6]. Although single-chain antibody binders with three CDR loops designed *in silico* were validated *in vitro*, the likelihood of generating effective binder sequences was estimated to be low, at less than 1%.

Moreover, the affinity of the resulting binders remained weak. The recent advancements introduced by AlphaFold 3 [7] are expected to further advance *de novo* antibody design.

The CDR-H3 loop (Complementarity Determining Region-Heavy chain loop 3, referred to as the H3 loop for simplicity) of an antibody plays a crucial role in target recognition and specificity due to its high sequence and structural variability. It has become a key focus in structure prediction, as current methods can accurately predict the overall antibody framework and the structures of the other five CDR loops, owing to the extensive data available from sequence and structure databases. The reported H3 loop structure prediction accuracy is typically above 2 Å [8, 9]. Here we show that the H3 loop structure can be predicted with nearly 1 Å accuracy, reaching atomic precision, when the approximate structure and orientation of the binding protein are provided.

Evaluation of protein structure prediction methods requires ground-truth experimental structures. *In silico* testing of a structure prediction method involves measuring the difference between the predicted and the ground-truth structures. However, for protein design tasks, a single ground-truth solution does not exist and numerous alternative solutions may be possible. An approximate way of *in silico* evaluation is performed by checking the consistency of the designed structure and the structure predicted from the designed sequence [2], which we call the *in silico* design success rate. We demonstrate a correlation between antibody loop structure prediction accuracy and the success rate of *in silico* target-binding antibody loop design based on evaluations of the two versions of our models. This underscores the important concept that accurate antibody sequence design requires accurate antibody structure prediction.

In this work, we also performed *in vitro* evaluation of our antibody binder design technique through wet-lab synthesis and binding measurement of the designed antibodies. An older version of our method (GaluxDesign™ v1) demonstrated a high *in vitro* success rate for zero-shot antibody loop design, achieving 13.2% compared to the previously reported rate of 1.8% [10] for the redesign of the three HCDR (Heavy chain CDR) loops of a HER2 (human epidermal growth factor receptor 2) antibody. A more recent version of our design method (GaluxDesign™ v2), which shows higher loop structure prediction accuracy and *in silico* loop design success rate, has shown further improvement in antibody loop design tasks targeting three additional therapeutic targets: human PD-L1 (programmed cell death 1 ligand 1), human PD-1 (programmed cell death protein 1), and human EGFR (epidermal growth factor receptor), with some of the designed antibodies showing sub-nanomolar affinity. For the EGFR case, antibodies with binding specificity to a mutant target (S468R) were designed *in silico* and confirmed *in vitro*. These results from antibody loop design experiments show promise for the next-stage design: *de novo* design of full antibody sequences to bind specified epitopes.

## 2 Results and Discussion

### 2.1 Performance of antibody H3 loop structure prediction

Current structure prediction methods exhibit lower accuracy in predicting antibody H3 loop structures compared to other protein structures. Figure 1 presents a performance comparison of available prediction methods for the H3 loop structure prediction task within an antibody–protein complex structure, with prediction error measured as RMSD (RootMean-Squared Deviation) of the C_*α*_ coordinates in the H3 loop. The recent Galux structure prediction model (v2) achieved the best performance of 1.4 Å RMSD on a newly compiled, time-separated set, the GAbD (Galux Antibody Design) set, created for an objective comparison of recently developed deep learning methods [9, 11, 12]. Detailed procedures for running the compared methods are provided in Section 4.2. For example, AlphaFold 2.3 [11] was modified to take an antibody–protein complex structure as a structural template with interchain features, instead of using MSA (Multiple Sequence Alignment), to measure antibody loop structure prediction performance without being affected by the prediction performance of antibody–protein orientations. ABlooper and ImmuneBuilder were also run in-house, as detailed in Section 4.2.

**Figure 1:**
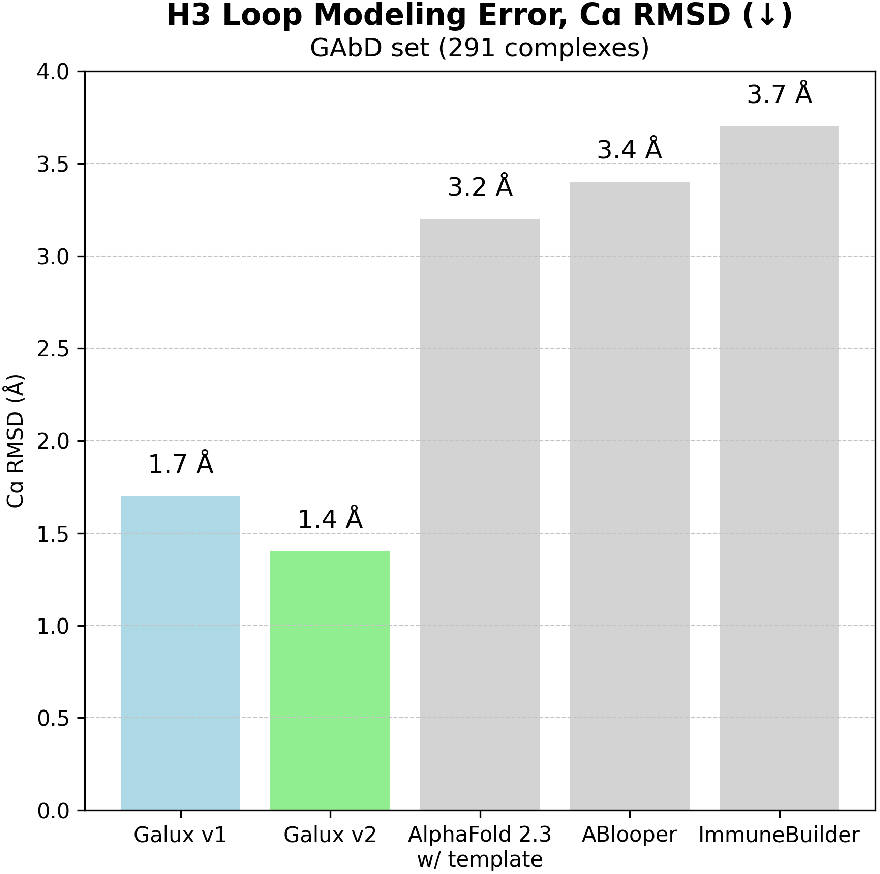
Mean H3 loop structure prediction error, measured in C_*α*_-RMSD, for five structure prediction methods on a set of 291 antibody H3 loops whose PDB structures were released after January 1, 2023.

The GAbD set consists of 291 antibody–protein complexes with experimentally resolved structures released after January 1, 2023. Complexes with HCDR and LCDR (light-chain CDR) sequence identities exceeding 70% relative to previously released structures were excluded. This set is more up-to-date compared to previous test sets, such as the RAbD (Rosetta Antibody Design) set [13], which consists of 60 antibody–protein complexes released before 2019. The RAbD set includes the dataset used to train some of the compared methods. The average length of H3 loops in the GAbD set is 12.6 amino acids, while that of the RAbD set is 11.0. These factors make it more challenging to predict the H3 loop structures in the GAbD set. For example, Galux v1, Galux v2, AlphaFold 2.3 with template [11], ABlooper [9], and ImmuneBuilder [12] showed errors of 1.0 Å, 1.0 Å, 2.4 Å, 1.8 Å, and 0.9 Å RMSD, respectively, on the RAbD set, but errors of 1.7 Å, 1.4 Å, 3.2 Å, 3.4 Å, and 3.7 Å, respectively, on the GAbD set.

Finally, it should be noted that AlphaFold 2.3 and ImmuneBuilder were not necessarily optimized for the specific task of H3 loop modeling, and their performance after potential fine-tuning for this task remains unknown.

### 2.2 *In silico* performance of antibody loop design

The performance of available protein design methods for the antibody loop sequence design task is illustrated in Figure 2. This task involved redesigning sequences of six antibody loops for a specified antibody–protein orientation across 291 antibody–protein complexes in the GAbD set. In Figure 2, loop design success rates are presented using two metrics: the structure recovery rate and the G-pass rate. The structure recovery rate measures the percentage of cases, out of 20 designs for a given antibody–protein orientation, where the designed H3 loop structures deviate by less than 2 Å in C_*α*_-RMSD from the experimentally resolved loop structure of the same length and orientation. The newly developed G-pass rate, which incorporates confidence and consistency scores from the Galux models, has proven to be a more stringent measure than the structure recovery rate.

**Figure 2:**
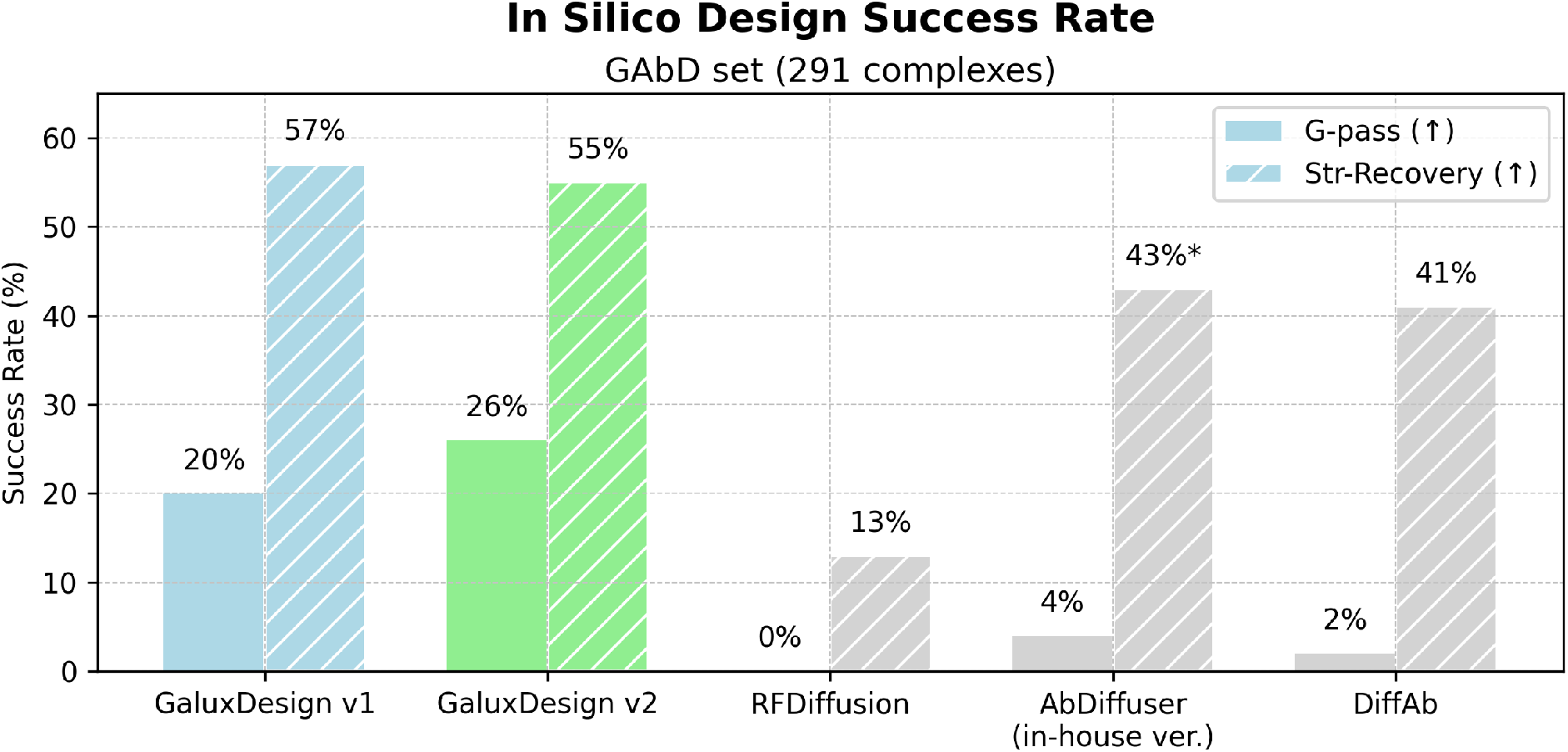
Comparison of the *in silico* performance of available protein design methods for the antibody loop sequence design task, evaluated using the structure recovery (Str-Recovery) rate and the G-pass rate derived from Galux structure prediction model (v2). To evaluate the structure recovery rate for AbDiffuser, only cases where the generated loops matched the reference in length (39% of the GAbD set) were included.

GaluxDesign v1 and GaluxDesign v2 show similar structure recovery rates (57% and 55%, respectively), but the G-pass rate provides better differentiation, highlighting the superior performance of GaluxDesign v2 (26%) compared to GaluxDesign v1 (20%). The other methods compared—RFdiffusion [2], AbDiffuser [14], and DiffAb [15]—demonstrated significantly lower success rates than the GaluxDesign methods in both structure recovery and G-pass rates. The high *in silico* antibody CDR loop design success rates of GaluxDesign models are attributed to leveraging the strong CDR loop structure prediction capabilities of Galux structure prediction models through the use of effective Galux generator models.

Although a fine-tuned RFdiffusion model specifically for antibody design was reported [6], a performance test of this version was not conducted due to program unavailability. Detailed procedures for running these design methods are provided in Section 4.3.

### 2.3 Performance of antibody binder/non-binder nation discrimi-

A large number of sequences can be generated *in silico* to identify antibodies targeting a specific epitope on a protein. However, a small fraction of these potential binder sequences has to be selected for *in vitro* wet-lab evaluation, making the scoring step crucial. In Figure 3, the ability of structure prediction models to discriminate binder sequences from non-binder sequences is evaluated using AUROC (Area Under Receiver Operating Characteristic curve) on the HER2-Trastuzumab mutant library, consisting of approximately 34,000 sequences, 26% of which are binders [16]. Galux scoring, based on the G-pass rate, outperforms AlphaFold 2.3 scoring based on PAE (Predicted Aligned Error) (AUROC = 0.529), with Galux v2 demonstrating the best performance (AUROC = 0.758). Publicly available binder/non-binder data essential for developing binder sequence scoring are scarce, and additional experimental data could significantly advance the scoring methods.

**Figure 3:**
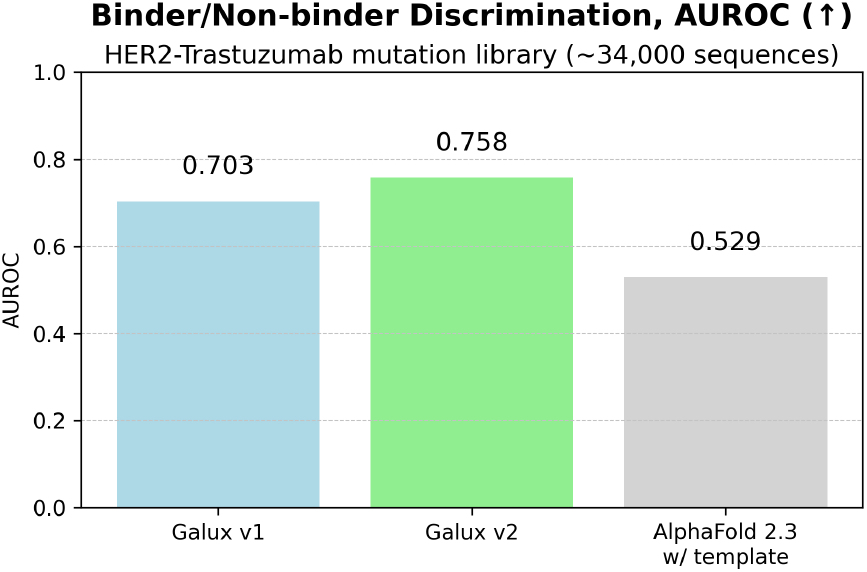
Performance comparison of binder scoring methods using confidence estimates from three structure prediction models on the HER2-Trastuzumab binder/non-binder set.

### 2.4 *In vitro* evaluation of antibody CDR loop design

A standard *in vitro* evaluation protocol for assessing the performance of *in silico* antibody CDR loop design has yet to be established. The performance of *de novo* antibody designs reported in the RFdiffusion-Ab study [6], validated *in vitro* for 3CDR loop designs in VHH (nanobody) frameworks targeting multiple proteins, is estimated to have a success rate of less than 1%. The success rate was calculated as the percentage of sequences showing detectable binding in *in vitro* binding assays. The success rate of less than 1% for RFdiffusion-Ab includes binders with very low affinity (K_*d*_ in the micromolar range).

A team from Absci reported antibody design results for a more focused task: zero-shot design of the three HCDR loops on a known antibody framework, specifically the HER2 antibody Trastuzumab [10]. The design algorithm was not informed by any target-specific information, such as known binder sequences for HER2, during training. A success rate of 1.8% was achieved for the zero-shot design of the three HCDR loops for the HER2 antibody [10].

Given the success rates in both reports, largescale *in vitro* tests using library display techniques with single-chain antibody formats (VHH or scFv) have been employed to evaluate the binding of the designed sequences.

Using the two GaluxDesign models (v1 and v2), we conducted the same type of zero-shot design experiments as the Absci team, designing CDR loop sequences within known antibody frameworks for four targets: HER2, PD-1, PD-L1, and EGFR. For EGFR, the CDR design was performed on the S468R mutant, even though the framework antibody is not a specific binder to this mutant. No known sequence or structural information was provided for the designed CDR loops. The target–antibody orientations were based on PDB structures (1N8Z, 5B8C, 5XXY, and 5SX4, respectively).

The *in vitro* evaluation results of the designed sequences, summarized in Table 1, show significantly improved performance compared to previous reports. Our team conducted two types of *in vitro* evaluations: one involved screening individually expressed antibodies in the IgG1 format (testing a smaller set of sequence designs for six CDR loops, approximately 50–60), while the other used yeast display in the scFv format (testing a larger set of sequence designs, around 10,000, for three HCDR loops). Details of the binding evaluation, including the criteria for defining binders in both methods, are provided in Section 4.5. In general, we classified antibodies with estimated binding affinities stronger than 100 nM as binders.

**Table 1:**
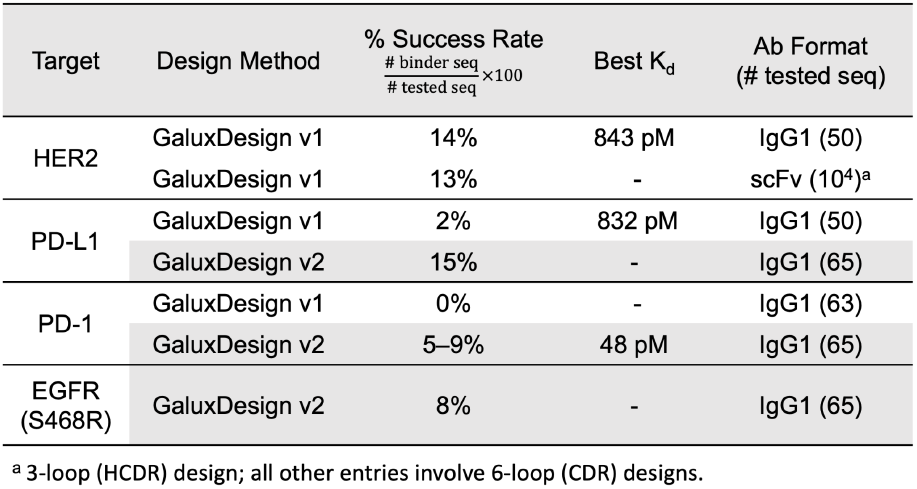
*In vitro* success rates, reflecting binders with estimated affinities tighter than 100 nM, for the design of target-binding antibody CDR loops using GaluxDesign models across four targets. The best binding affinity among the measured designs is also reported when available.

The earlier design method, GaluxDesign v1, was tested on a 3-loop HCDR design for a HER2 antibody, allowing for comparison with the results of the Absci team. Our success rate of 13%, determined using yeast display, is significantly higher than the previously reported rate of 1.8%. GaluxDesign v1 was also tested on the 6-loop CDR design using known antibody frameworks for HER2, PD-L1, and PD-1, yielding success rates of 14%, 2%, and 0%, respectively, based on an in-house protocol for binding evaluation of individually expressed IgG1 antibodies. The higher success rate for HER2-targeting antibody design than the other targets suggests that the provided antibody orientation information played a more significant role in the HER2 case. Antibodies with sub-nanomolar affinities were obtained for HER2 and PD-L1, as detailed in the next section.

The more recent version, GaluxDesign v2, was tested on the 6-loop design of PD-L1, PD-1, and EGFR (S468R mutant) antibodies, resulting in success rates of 15%, 5–9%, and 8%, respectively. The increased success rates for PD-L1 (from 2% to 15%) and PD-1 (from 0% to 5–9%) compared to v1 are consistent with the *in silico* performance evaluations for loop design and binder/non-binder discrimination using the G-pass rate. The design results for the newly tested target, the S468R mutant of EGFR, are also noteworthy, leading to mutant-specific antibodies as detailed in the next section.

Beyond the CDR loop design experiments within a given antibody orientation, *de novo* design experiments without orientation information is currently underway using further improved GaluxDesign models.

### 2.5 Sequence diversity and novelty, target binding affinity, and mutant specificity of the designed antibodies

A subset of antibody binders obtained from the initial *in vitro* screening was subjected to further analysis, as summarized in Table 2. The loop length and sequence identity of the designed CDR loops relative to the reference antibody show the greatest variability in the H3 loops, followed by the L3 loops across all four targets. The H3 loops of the binder antibodies in the table exhibit relatively high sequence diversity and novelty, with sequence identity to the reference ranging from 25% to 50%.

**Table 2:**
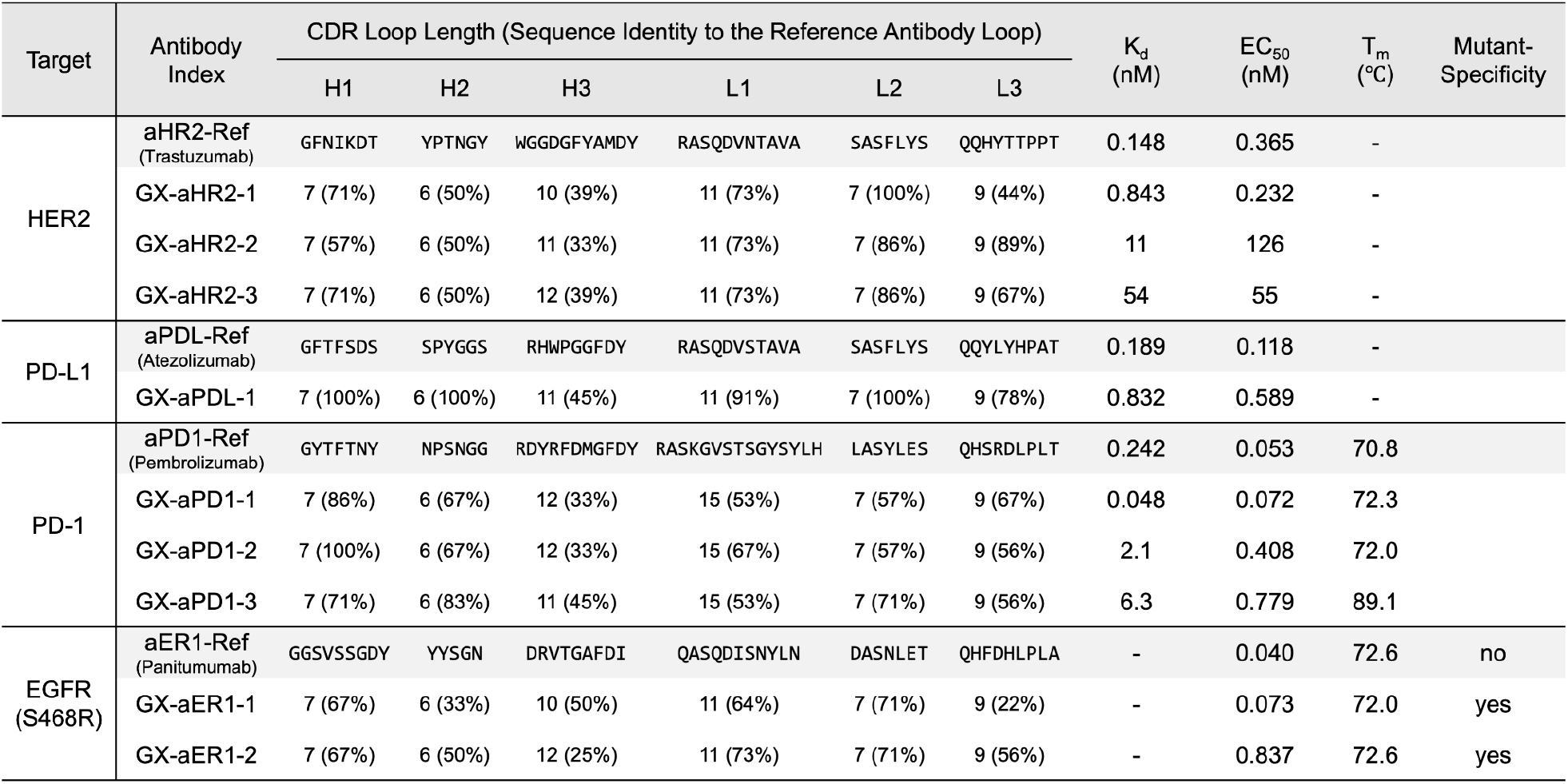
Sequence length and identity to reference, binding affinity (measured by K_*d*_ and EC_50_), melting temperature (T_*m*_), and mutant specificity (EGFR case) for the sampled designed antibodies.

Binding affinity to the respective target was measured for selected antibodies using ELISA and BLI, as shown in Figure 4. For all four targets, the best binding affinities achieved were in the subnanomolar range, with a few cases reaching two-digit picomolar affinity, as shown in Table 2. The K_*d*_ value for the designed antibodies targeting EGFR-S468R is still under investigation.

**Figure 4:**
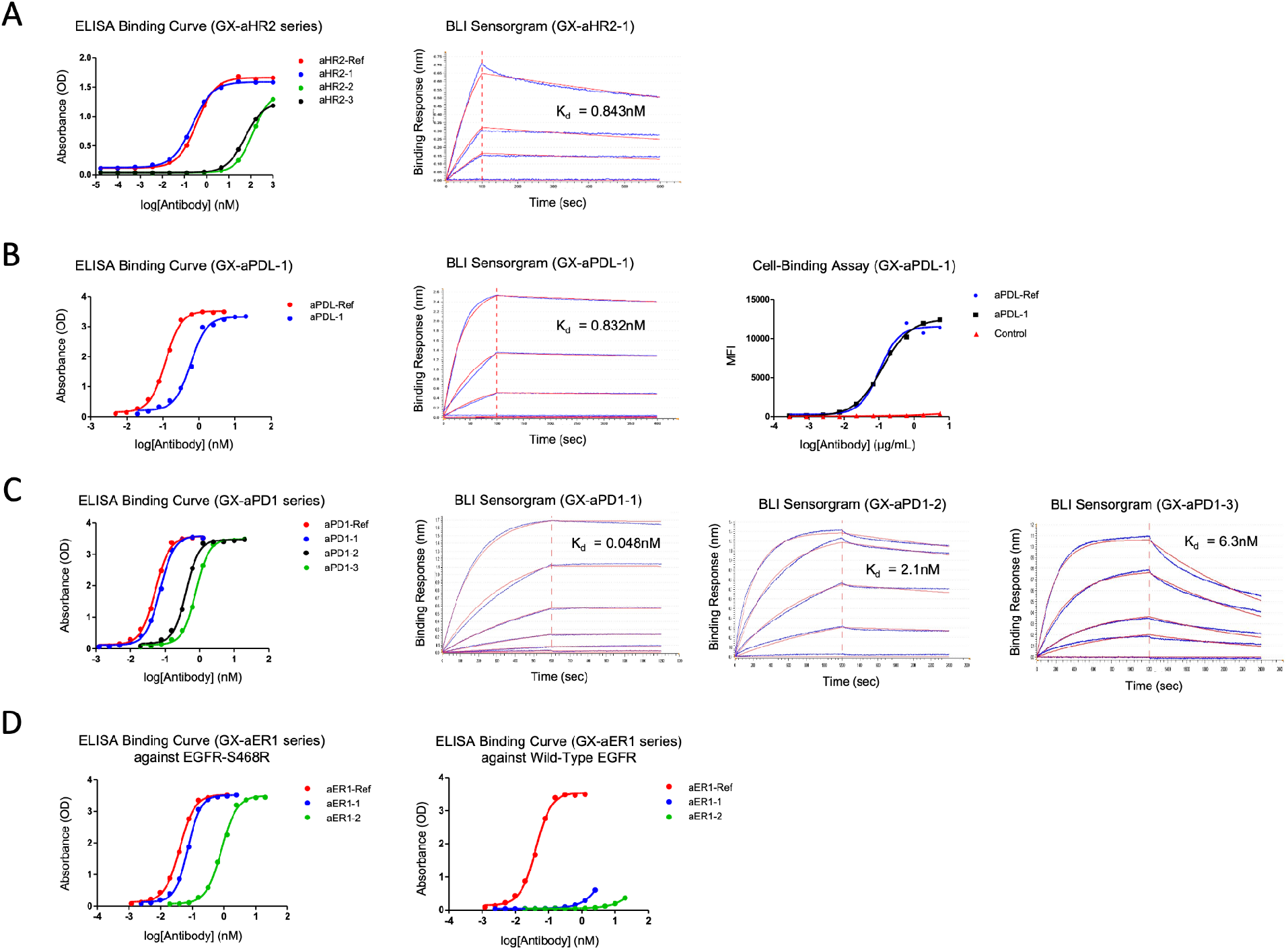
*In vitro* binding affinity measurements by ELISA (A, B, C, and D) and BLI (A, B, and C), cell binding assay (B), and mutant specificity test (D) for selected antibody binders. (A) GX-aHR2 antibodies targeting HER2: ELISA curve of GX-aHR2-1 (left) and BLI sensorgram of the GX-aHR2 series (right). (B) GX-aPDL-1 antibody targeting PD-L1: ELISA curve (left), BLI sensorgram (middle), and cell-binding assay (right) for GX-aPDL-1. (C) GX-aPD1 antibodies targeting PD-1: ELISA curves (left) and BLI sensorgrams (right three plots) for the GX-aPD1 series. (D) GX-aER1 antibodies targeting EGFR-S468R: ELISA curves against EGFR-S468R (left) and wild-type EGFR (right) for the GX-aER1 series.

For the GX-aPDL-1 antibody designed to target PD-L1, a cell-binding assay demonstrated a strong binding signal comparable to that of the reference. (Figure 4B, right).

The antibodies GX-aER1-1 and GX-aER1-2 were designed to target the S468R mutant of EGFR. Binding specificity to the mutant was confirmed, as these antibodies showed significantly weaker binding to wild-type EGFR compared to EGFR-S468R (Figure 4D). These results demonstrate the capability of GaluxDesign v2 to design antibodies that bind to intended epitopes with precise target specificity.

To further confirm that the designed antibodies target the intended epitopes, a competition binding assay was performed using BLI with reference antibodies against PD-L1 and HER2. The results (data not shown) revealed that GX-aPDL-1, designed to target PD-L1, could not bind PD-L1 simultaneously with Atezolizumab, the reference PD-L1 antibody. Likewise, GX-aHR2-1, designed to target HER2, could not bind HER2 simultaneously with Trastuzumab, the reference HER2 antibody. These findings indicate that the designed antibodies occupy the same epitopes as the reference antibodies.

Examples of the predicted structures of experimentally confirmed binder antibodies in complex with their targets are shown in Figure 5. The designed antibodies GX-aHR2-1, GX-aHR2-2, and GX-aHR2-3, which target HER2, exhibit structural diversity, including variability in H3 loop length (Table 2) and protein–antibody orientation (Figure 5A), likely stemming from CDR loop sequence diversity. The GX-aPDL-1 antibody features a compact binding interface with its target protein, PD-L1, characterized by favorable hydrophobic and polar interactions, with atomic-level side-chain interactions involving loop residues (Figure 5B).

**Figure 5:**
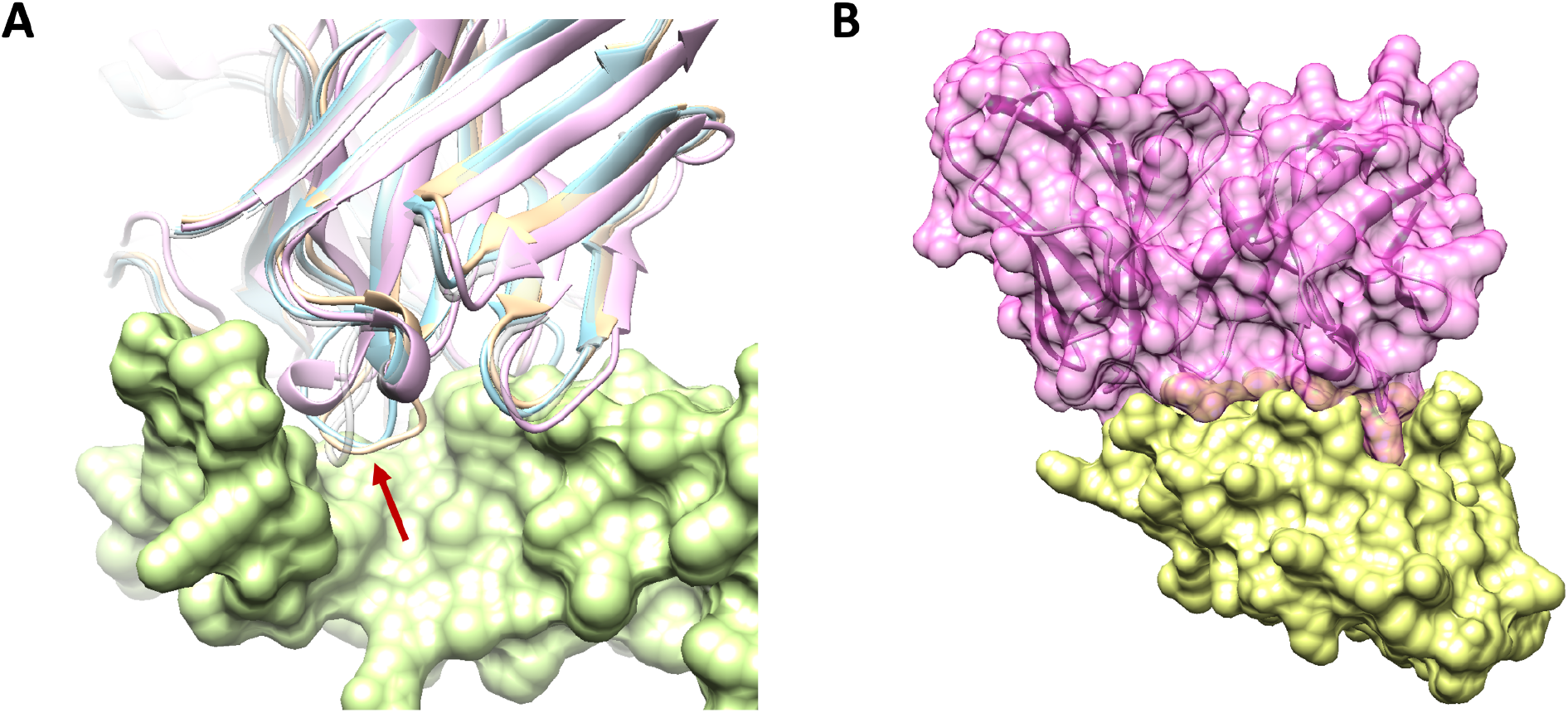
Predicted structures of selected designed antibodies in complex with their targets. (A) The backbone structures of three antibodies (tan, blue, and pink) designed to target HER2 (green), compared to the reference antibody (transparent gray), demonstrate structural diversity in the H3 loop (pointed by the red arrow). (B) A designed antibody (magenta) in complex with its target, PD-L1 (lime), exhibits high shape complementarity between the two.

## 3 Conclusion

In this study, we demonstrated the crucial role of accurate antibody loop structure prediction in enabling effective zero-shot design of target-binding antibody loops. Our findings reveal that reliable loop structure prediction underpins robust loop design methodology. Specifically, the latest version of our prediction and design method, Galux v2, showed significant improvements in both the accuracy of antibody H3 loop structure predictions and the success rate of *in silico* loop designs. The effectiveness of this design method was further confirmed by experimental binding measurements. These advancements underline the potential of our approach for designing antibodies with high affinity and specificity, which are essential for therapeutic and industrial applications.

Looking ahead, advancing the design methodology as outlined in the development strategy could open new avenues for the *de novo* design of full antibody sequences targeting specific epitopes. The difficulty of *de novo* design will depend on the extent of structural changes induced by antibody–target binding, which can range from sidechain conformational adjustments to larger backbone rearrangements. Numerous potential therapeutic targets involve varying degrees of conformational change upon interaction with partner molecules. We believe that successful efforts in this area will solidify high-performance structure-based design as a foundational strategy in the next generation of antibody engineering.

## 4 Methods

### 4.1 Database curation for training Galux models and benchmarking methods

#### Protein structure database used for training the Galux models

All PDB protein structures [17] released before June 13, 2024, were collected, excluding structures whose HCDR and LCDR sequences have >70% sequence identities to entries in the Rosetta Antibody Design (RAbD) set [13], as well as antibody structures targeting the proteins HER2 and PD-L1 tested in this study. Structures released before December 31, 2022, were used for training and validating the Galux models. A time-separated antibody–protein complex set, named as GAbD (Galux Antibody Design) set, consisting of the structures released after January 1, 2023, excluding those with HCDR and LCDR sequence identities greater than 70% relative to the CDR sequences in the training and validation sets, was used for benchmarking predictive and generative models.

Chothia numbering [18], as implemented by SAbDab [19] or ANARCI [20], was used for antibody CDR designation. Among those in the RAbD set, six entries (2XQY, 1A14, 5EN2, 1FE8, and 3S35, which contain carbohydrates near the antibody–protein interface, and 5GGS, which is similar to 5B8C) were excluded from the benchmark.

#### Binder/Non-binder Database

The binding data for a HER2-Trastuzumab mutant library, as reported by [16], were utilized for the binder/nonbinder discrimination benchmark. This dataset includes 8,955 unique binder sequences and 25,191 unique non-binder sequences, after excluding those designated as both binder and non-binder. Each sequence is identical to Trastuzumab except for variations in the H3 loop sequence.

### 4.2 Measuring the performance of available structure prediction methods in antibody H3 loop structure prediction

In the antibody H3 loop structure prediction task, the input consists of the antibody–protein complex structure, excluding the coordinates of the six CDR loop regions. Prediction methods then generate the coordinates of the antibody loops within the provided structural environment. The prediction accuracy of the H3 loop structure was measured by C_*α*_-RMSD of the H3 loop structure after superimposing the non-CDR antibody framework structure onto the corresponding reference, ground-truth antibody structure.

The three compared methods for the H3 loop structure prediction task were run in-house as follows. First, “AlphaFold 2.3 w/ template” uses AlphaFold-Multimer [11] modified to take an antibody–protein structure as a structural template with inter-chain features instead of using MSA. This approach focuses on measuring antibody loop structure prediction performance without being affected by other parts of the antibody–protein complex structure prediction. Among the five model structures generated from the five parameter sets of AlphaFold v2.3, the highest-ranked model (as provided by AlphaFold v2.3) was selected for performance evaluation. Second, “ABlooper”, an end-to-end deep learning-based CDR loop structure prediction method [9], was run using the Chothia numbering scheme to designate the loop regions to model instead of the IMGT scheme. Lastly, “ImmuneBuilder”, which predicts the whole antibody structure from its sequence [12], was also run in-house using the standard protocol.

### 4.3 *In silico* performance measurement of antibody loop design

The antibody loop design was conducted using the antibody–protein complex structure as input, excluding the sequences and coordinates of the six CDR loops. The coordinates and sequences of the CDR loops of the same lengths were then generated within the given structural environment. The different design methods were compared by generating 20 antibody loop sequences for each entry of the GAbD set.

Two performance metrics, G-pass and Str-Recovery (structure recovery), were used to assess the effectiveness of the design methods. The G-pass rate evaluates the effectiveness of each sequence design based on the confidence of structure prediction and the consistency between the designed structure and the structure predicted from the designed sequence, as estimated by Galux structure prediction model v2. The structure recovery rate, measures the percentage of cases where the minimum C_*α*_RMSD between the designed H3 loop structure and the corresponding crystal loop structure is less than 2 Å.

The three design methods, RFdiffusion, AbDiffuser, and DiffAb, were run as follows. First, the CDR loop design labeled as “RFdiffusion” in Figure 2 was conducted by generating the CDR backbone structure with RFdiffusion [2], followed by sequence prediction on the designed backbone structure using ProteinMPNN [21]. Next, AbDiffuser, an antibody-specific diffusion model for simultaneous sequence and structure generation [14], was reproduced in-house (“AbDiffuser in-house ver.”) for performance comparison. Since the length of the loop generated by AbDiffuser could not be controlled, only designs with the same loop lengths as the template—comprising 39% of cases—were used to evaluate the structure recovery rate. Therefore, this value cannot be directly compared to those of the other methods. Finally, “DiffAb”, an antibody-specific diffusion model for sequence-structure codesign [15], was evaluated using the standard protocol.

### 4.4 *In silico* performance measurement of antibody binder/nonbinder discrimination task

In the antibody binder/non-binder discrimination task, the input consists of the predicted antibody H3 loop structure in the environment of the reference complex structure (PDB entry 1N8Z). Scoring methods are expected to predict a measure of binding affinity, considering the provided antibody–protein complex structure. AUROC is calculated for each prediction result after obtaining all score values for binder and non-binder sequences in the database described in Section 4.1.

In this study, we compared the confidence score of the Galux models with the interface PAE score (calculated as the average PAE value of residue pairs involved in antibody–protein interactions) of “AlphaFold 2.3 w/ template” as scoring metrics, as such confidence scores for protein–protein interfaces are often used as proxies for binding affinity [22]. However, the physical chemistry of binding affinity is far more complex, involving interactions with solvent molecules that must be accounted for as ensemble averages.

### 4.5 *In vitro* evaluation of six CDR loop design

#### Computational design

For four target proteins, the six CDR loops were designed using GaluxDesign models, based on the provided structure of the antibody–protein complex (HER2, PD-L1, PD-1, and EGFR-S468R: PDB entry 1N8Z[23], 5B8C[24], 5XXY[25], and 5SX4[26], respectively), excluding the CDR regions. During the training and validation of the Galux methods, as well as during the design process, the sequence and structural information of the CDR regions were not utilized. The number of designed antibodies selected for *in vitro* validation is listed in Table 1.

#### Initial screening

To construct antibodies in the IgG1 format, we synthesized each DNA as a gBlock™ from IDT and performed cloning. The gene was inserted into the pcDNA™3.4 plasmid and transfected into Expi293F™ cells. The concentration of the expressed antibody in the supernatant was measured by BLI before purification. To rapidly screen many antibodies, we performed ELISA assay twice at different concentrations (10 *µ*g/mL and 0.1 *µ*g/mL) of antibodies in the supernatant. Each test was performed in triplicate. The optical density (OD) value cutoff was determined by calibrating with experimentally determined K_*d*_ values of selected antibodies. The specificity of the antibodies against off-target proteins was also assessed in this stage.

#### Binding assay

From the initially screened antibodies, a subset was selected for purification and affinity measurement. Binding affinity was evaluated using EC_50_ via ELISA and K_*d*_ via BLI. A competition binding assay was also conducted using BLI. For GX-aHR2-1, Biotin-HER2 was first loaded, followed by 250 nM of Trastuzumab (which shares binding site with GX-aHR2-1, the fourth domain of HER2) or Pertuzumab (which binds to different interface, the second domain of HER2), and then 250 nM of GX-aHR2-1, with the order of addition reversed in a separate experiment. For GXaPDL-1, Biotin-PD-L1 was first loaded, followed by 250 nM of Atezolizumab, and then 250 nM of GX-aPDL-1. In the cell binding assay, a PD-L1 overexpressing stable cell line was constructed using HEK293Ta cells, and the cell binding test was performed by binding individual antibodies and measuring the results with a cell analyzer.

### 4.6 *In vitro* evaluation of three HCDR loop design

#### Computational design

The three heavy chain CDR loops of Trastuzumab were re-designed using GaluxDesign v1, based on the the available structure of HER2-Trastuzumab complex (PDB entry 1N8Z), excluding the three heavy chain CDR regions. A total of 10,000 antibodies were designed. During the training and validation of the Galux models, as well as throughout the design process, the sequence and structural information of the CDR regions were not utilized.

#### Library-based binder evaluation

To construct the scFv antibody library, we synthesized a DNA oligo pool from Twist Bioscience and performed cloning. The gene was inserted into the pYD1 yeast plasmid, transformed into *E. coli* for cloning, and subsequently into EBY100 for yeast expression.

After constructing the scFv library, we conducted yeast display with two rounds of biopanning using the Sony SH800S sorter. The on-target and offtarget (PD-L1) binding signals of the screened pool demonstrated the specific binding properties of the designed antibodies. Next-generation sequencing (NGS) of the screened pool identified 1,320 designed sequences enriched across two rounds of screening. From these, 30 individual antibodies were randomly selected and confirmed to specifically bind to HER2 with affinities tighter than approximately 100 nM (data not shown). As a result, the design success rate was estimated to be 13.2% (1,320/10,000 × 100%).

## 5 Author Contributions

T. P., C. S., J. W., and J. Y. conceived the study and coordinated the collaborations. W. B., J. G., J. K., S. K., Dohoon L., Dongjin L., J. Lee, Y. Lee, J. Lim, J. Nam, J. Noh, S. S., J. W., H. W., H. Y., and J. Y. developed the predictive and generative AI methods for protein structure prediction and design. Y. B., Y.-A. C., E. S. L., M. S. L., Sangchoon L., Soyul L., S. J. L., J. Y. M., H. O., S.-Y. P., M. Y. S., and M. J. Y. conducted the wet-lab validations. Sangchoon L., C. S., M. Y. S., and J. W. wrote the initial manuscript draft. All authors contributed to the final version.

